# Lipid chaperone LBP-8 coordinates with nuclear factors to promote longevity in *Caenorhabditis elegans*

**DOI:** 10.1101/2021.09.09.459489

**Authors:** Jonathon Duffy, Qinghao Zhang, Sung Y. Jung, Meng C. Wang

## Abstract

Eukaryotic cells are composed of a variety of organelles. Their coordination plays crucial roles in cellular homeostasis and organism longevity and is mediated by proteins with the ability to transport between different organelles. In *Caenorhabditis elegans*, LBP-8 is a pro-longevity lipid chaperone that can localize to both lysosomes and the nucleus. Here we profiled LBP-8’s binding partners using immunoprecipitation and mass spectrometry. From the 45 identified candidates, we discovered four nuclear factors that are required for the LBP-8-induced longevity. Among them, RPC-2, an RNA Polymerase III core subunit, is also necessary for the nuclear localization of LBP-8. Moreover, we have screened nuclear transport machinery components, and revealed the requirement of the nuclear import, not export, for the LBP-8 longevity effects. Together, these results suggest that the lipid chaperone LBP-8 relies on specific nuclear factors to retain in the nucleus and regulate longevity.

## Introduction

Lipids are key bioactive molecules that play a number of important roles, including supplying and storing energy, providing compartmentalization to allow specialized functions within a cell, and modulating signal transduction (1). As signaling molecules, specific lipids can directly function as ligands for receptors such as G-protein coupled receptors and nuclear hormone receptors (NHRs) and as secondary messengers, and they can also modulate signaling mechanisms through altering protein configuration and localization in the membrane (1).

Lipids are not hydrophilic, which makes it difficult for them to diffuse freely through the aqueous environment. One group of lipid chaperones that facilitate lipid transportation and signaling effects are the intracellular lipid binding proteins (iLBPs), which include the Cellular Retinol Binding Proteins (CRBPs), Cellular Retinoic Acid Binding Proteins (CRABPs), and Fatty Acid Binding Proteins (FABPs) (2). These small (12-20kDa) proteins display a characteristic structure of a beta barrel with an alpha-helix lid, show a promiscuous ligand binding capability, and have a partially overlapping tissue expression pattern (3,4). Among them, several FABPs have been shown to localize to the nucleus (3). For example, FABP4 and FABP5 both have a nuclear localization sequence (NLS) (5,6), and they cooperate with peroxisome proliferator-activated receptors (PPARs) to regulate metabolic and inflammatory processes (7). The nuclear retention of FABP4 relies on the exportin system (5), and specific lipid binding influences the nuclear translocation of FABP5 (6). Dependent on the identity of the bound lipid ligand, the NLS of FABP5 can be either stabilized or destabilized (6). Interestingly, FABP4/5 deficiency in mice attenuates age-associated weight gain, glucose intolerance, and hepatosteatosis, but shows no protection against age-associated declines in cardiac, muscular or cognitive functions (8). In addition, despite the improvement of metabolic fitness, the FABP4/5 deficient mice exhibit no changes in their lifespan (8). For the other eight FABPs in mammals, their effects on lifespan and healthspan remain untested.

In *Caenorhabditis elegans*, there are 9 FABP encoding genes, termed *lbp*’s. LBP-8 has been discovered as the first pro-longevity lipid chaperone. *lbp-8* is specifically expressed in the intestine, the major fat storage tissue of *C. elegans*, and its proteins are detected both at the lysosome and in the nucleus (9). Upon the induction of a lysosomal acid lipase *lipl-4*, the *lbp-8* gene is transcriptionally up-regulated and the nuclear translocation of the LBP-8 protein is enhanced (9). The loss-of-function mutant of *lbp-8* shows no changes in lifespan, but partially suppresses the pro-longevity effect conferred by *lipl-4* overexpression (9). The overexpression of *lbp-8* prolongs lifespan and reduces fat storage via the upregulation of mitochondrial β-oxidation genes (10). Moreover, the LBP-8 protein has a high degree of structural similarity to its mammalian counterparts (11) and binds to several different fatty acids and a fatty acid derivative oleoylethanolamide (9,11). Importantly, the translocation of LBP-8 from the lysosome to the nucleus facilitates the activation of a NHR complex NHR-49/NHR-80 (9), which is crucial for the longevity-promoting effect (9). However, the molecular mechanism mediating the nuclear localization of LBP-8 remains unclear.

In this study, we have profiled the binding partners of LBP-8 through proteomic analysis, revealing an enrichment of nuclear proteins. Through a genetic screen of these binding partners and also factors involved in the nuclear transportation system, we identified specific proteins that mediate both the lifespan extending effect and the nuclear translocation of LBP-8. These findings reveal the regulatory mechanism by which LBP-8 promotes longevity through coordinating with specific nuclear factors.

## Results and Discussion

### Profile binding partners of pro-longevity lipid chaperone LBP-8

*lbp-8* is expressed at a low level in wild type conditions, which is induced upon lysosomal lipolysis to promote longevity (9). Constitutive expression of *lbp-8* on its own sufficiently prolongs lifespan (9). To understand the molecular mechanism by which LBP-8 regulates longevity, we have systemically searched for its protein binding partners through proteomic profiling. To this end, we generated a transgenic strain of *lbp-8* that expresses 3XFLAG fusion of LBP-8 under its endogenous promoter (*LBP-8::3xFLAG Tg)*. This transgenic strain is long-lived compared to the control (Figure 1A). We then performed immunoprecipitation (IP) with the anti-3XFLAG antibody in both the *LBP-8::3xFLAG Tg* strain and wild type (WT) worms, which is followed by mass spectrometry (MS) profiling of proteins (Supplemental Table 1). We first confirmed that the three independent replicates of IP-MS proteomic profiling exhibit a good degree of correlation (R=0.89-0.94) with each other (Figure 1B). Next, for each replicate, we calculated the fold change of each identified protein in MS between the *LBP-8::3xFLAG Tg* and WT samples and used the cut-off of at least 50-fold enrichment in *LBP-8::3XFLAG Tg*. There are seven candidates plus LBP-8 that are enriched in all three replicates, and 37 candidates that are enriched in two out of three replicates but not detected in either the *LBP-8::3XFLAG Tg* or WT worms in the third replicate (Figure 1C, Supplementary Table 1). Given that low abundant proteins might not be always detected by MS, we have included all of these 45 proteins as candidates (Table 1).

**Table 1:**
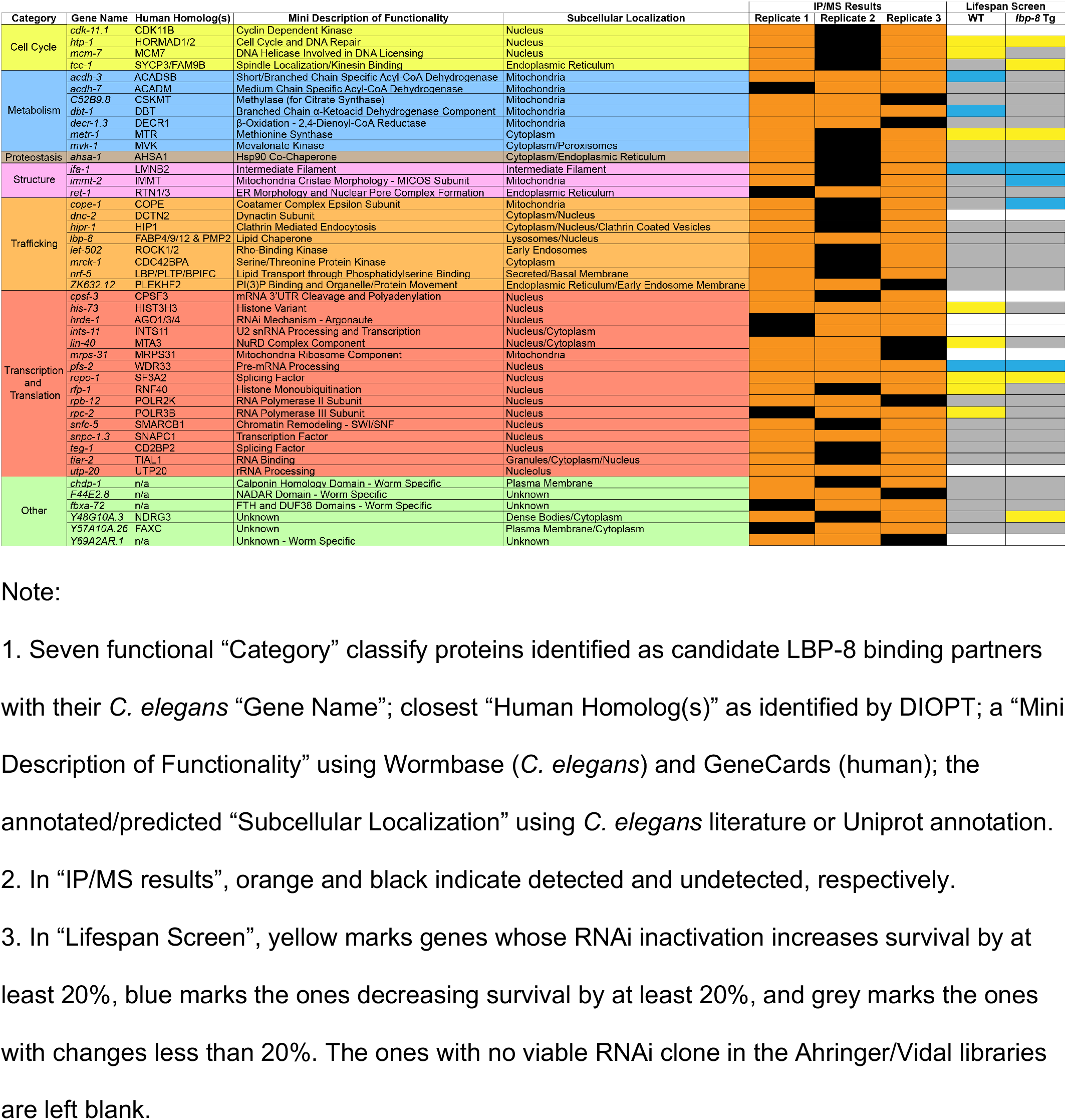
Summary of LBP-8 Binding Partners.

**Figure 1:**
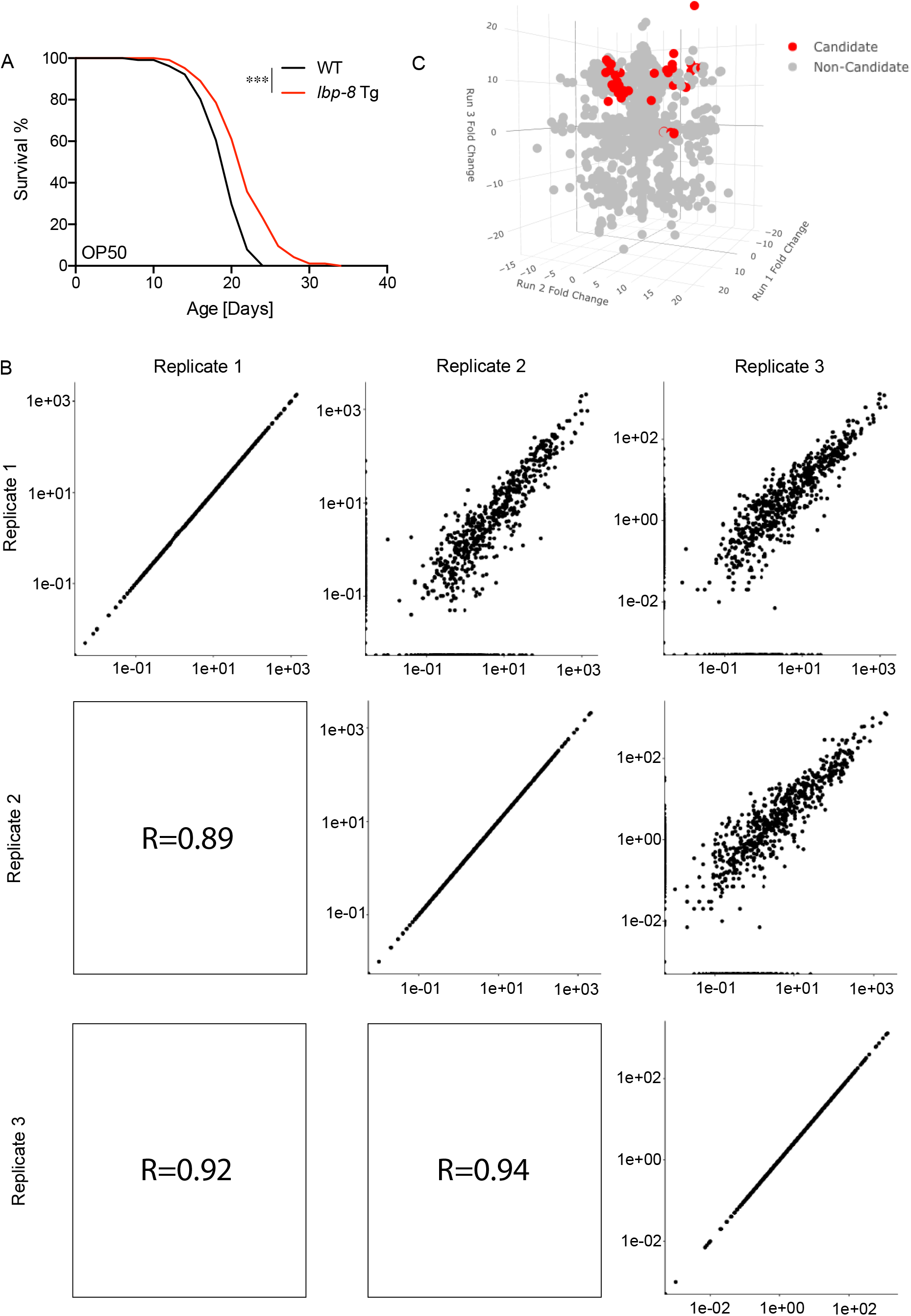
Proteomic Profiling of LBP-8::3xFLAG Binding Partners. (A) Overexpression of *LBP-8::3xFLAG* extends lifespan, *p<0*.*001 Tg vs. WT* by a log-rank test. (B) The correlation of the three replicates: in the upper right 3 panels, the iBAQ scores of each replicate are plotted for pairwise comparisons, and in the bottom left 3 panels, the Pearson correlation coefficients for each pairwise comparison are shown. (C) A 3D plot of the fold changes of proteins (*Tg* vs WT) for each replicate along a separate axis. Highlighted in red are the candidate binding partners.

Using DIOPT, we identified the closest mammalian homolog(s) of 41 of the 45 candidates (Table 1). Next, based on the identity of both worm proteins and mammalian homologs and their functions described and/or predicted on both WormBase and GeneCards, respectively, we categorized these candidates into seven functional groups: Cell Cycle, Metabolism, Proteostasis, Structure, Trafficking, Transcription and Translation, and Unknown Function (Table 1). In addition, we annotated the subcellular localization of each candidate using publications on the *C. elegans* proteins or Uniprot annotations of the human homologs. Among these binding partners, we found a noticeable enrichment of proteins annotated with either mitochondrial or nuclear localization, which is in line with altered mitochondrial physiology associated with the LBP-8-induced longevity (10) and the lipid chaperone function of LBP-8 to facilitate the nuclear delivery of lipid ligands (9).

### Elucidate lifespan regulatory effects of LBP-8 binding partners

To examine whether the binding partners of LBP-8 play a role in regulating longevity, we performed a screen using RNA interference (RNAi) of the encoding genes of these proteins in both the *LBP-8::3XFLAG Tg* and WT worms. To facilitate the screen, 5-fluoro-2’-deoxyuridine (FUDR) is administered to inhibit reproduction. On day 17 of adulthood, we assessed how many worms were still alive. Across the 4 replicates of the empty vector (EV) controls, the average percentage of WT worms alive was 38% and the average percentage of the *LBP-8::3XFLAG Tg* worms alive was 56%. To identify candidates that change the lifespan of either genotype, we used a cutoff of a 20% change in survival compared to the EV controls. Based on their effects, the LBP-8 binding partners were grouped into four categories (Figure 2A, Supplementary Table 2). Category 1 consists of *cope-1* and *immt-2* whose RNAi inactivation decreases the survival rate of the *LBP-8::3XFLAG Tg* strain but not WT. Both COPE-1 and IMMT-2 localize to mitochondria (12,13). Category 2 consists of *ifa-1* and *pfs-2* whose RNAi inactivation decreases the survival rate of both WT and the *LBP-8::3XFLAG Tg* worms, but the reduction extent is greater in the *LBP-8::3XFLAG Tg* worm than in WT. IFA-1 is a structural protein of intermediate filaments and PFS-2 is a nuclear protein (14,15). Category 3 consists of five genes, including *his-73, lin-40, mcm-7, rfp-1* and *rpc-2*, whose RNAi inactivation increases the survival rate of WT but causes no further increase in the *LBP-8::3XFLAG Tg* worms. All five proteins localize to the nucleus in *C. elegans* (16,17), or their homologs do in mammals (18). Category 4 consists of three genes, *htp-1, metr-1* and *repo-1*, whose RNAi inactivation increases the survival rate of both WT and the *LBP-8::3XFLAG Tg* worms. Both HTP-1 and REPO-1 are nuclear proteins (18,19). Thus, nuclear proteins are enriched in the candidates that affect lifespan.

**Figure 2:**
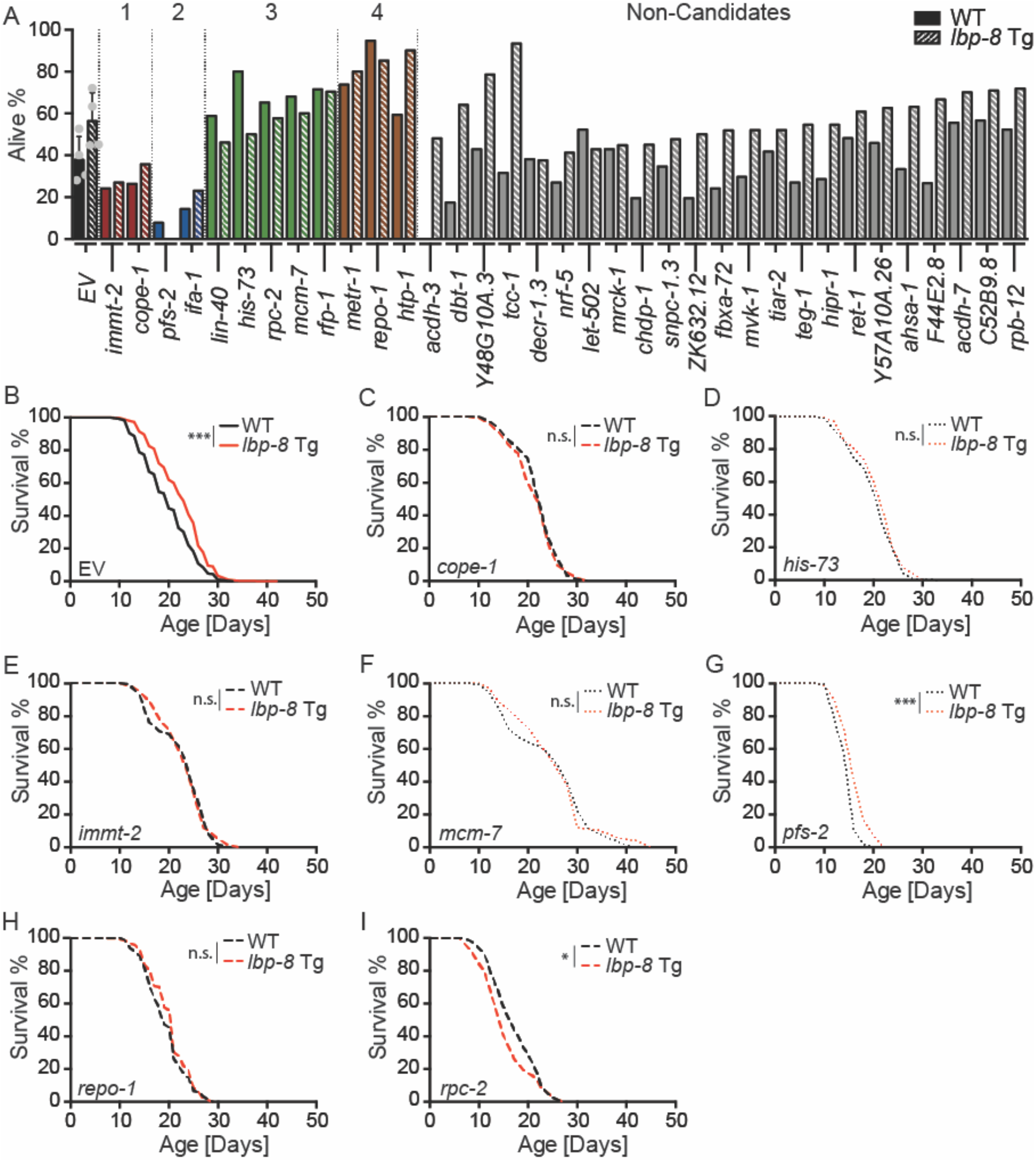
LBP-8 Binding Partners Regulate Longevity. (A) Summary of longevity regulating binding partners of LBP-8: Group 1 (Red) RNAi decreasing the survival of *LBP-8::3xFLAG Tg* but not WT worms; Group 2 (Blue) RNAi decreasing the survival of both; Group 3 (Green) increasing the survival of the WT but not the *LBP-8::3xFLAG Tg*; Group 4 (Brown) increasing the survival of both. Screening cutoff, >20% change in survival at day-17 adulthood. (B-I) Longitudinal lifespans of the *LBP-8::3XFLAG Tg* and WT worms with EV or RNAi inactivation. The *LBP-8::3XFLAG Tg* displays lifespan extension on the EV control (B). The *LBP-8::3XFLAG Tg* cannot extend lifespan with the RNAi inactivation of *cope-1* (C), *immt-2* (E), *repo-1* (H), or *rpc-2* (I) starting at the L1 stage; or with the RNAi inactivation of *his-73* (D) or *mcm-7* (F) starting the L4 stage. The RNAi inactivation of *pfs-2* starting at the L4 stage does not affect the lifespan extension conferred by *LBP-8::3XFLAG Tg* (G). *\*p<0*.*05, ***p<0*.*001 Tg vs. WT* by a log-rank test.

Next, we validated the effects of seven candidates (*cope-1, his-73, immt-2, mcm-7, pfs-2, repo-1* and *rpc-2*) on lifespan using longitudinal assays without FUDR. We found that except for *pfs-2*, RNAi inactivation of all the other candidates fully suppress the lifespan extension in the *LBP-8::3XFLAG Tg* worms (Figure 2 B-I). The mitochondrially localized proteins COPE-1 and IMMT-2 are the epsilon subunit of the coatamer complex and an inner mitochondrial structural protein, respectively. *his-73* encodes a histone variant, and *mcm-7* encodes a licensing factor involved in DNA replication. *rpc-2, pfs-2*, and *repo-1* encode components involved in RNA transcription and maturation, which are a core component of the RNA polymerase III complex, a polyadenylation factor, and a splicing factor, respectively. Together, these results confirm that the nucleus-localized binding proteins of LBP-8 are crucial for its pro-longevity effects.

#### RPC-2/RNA Pol III specifically regulates LBP-8 nuclear translocation and longevity

Next, to determine whether these lifespan-suppressing candidates affect the nuclear localization of LBP-8, we generated a transgenic strain expressing LBP-8 fused with GFP under the control of the *lbp-8*’s endogenous promoter and imaged the worms using confocal fluorescence microscopy. Consistently with our previously findings [48], we found that the LBP-8::GFP fusion protein was detected in the intestine, with the strongest expression in the first and last pairs of intestinal cells. At the subcellular level, the LBP-8::GFP fusion protein colocalizes with the RFP fusion of the lysosomal membrane protein LMP-1 (Figure 3A) and is also detected in the nucleus marked by the nuclear DNA dye Hoechst (Figure 3B).

**Figure 3:**
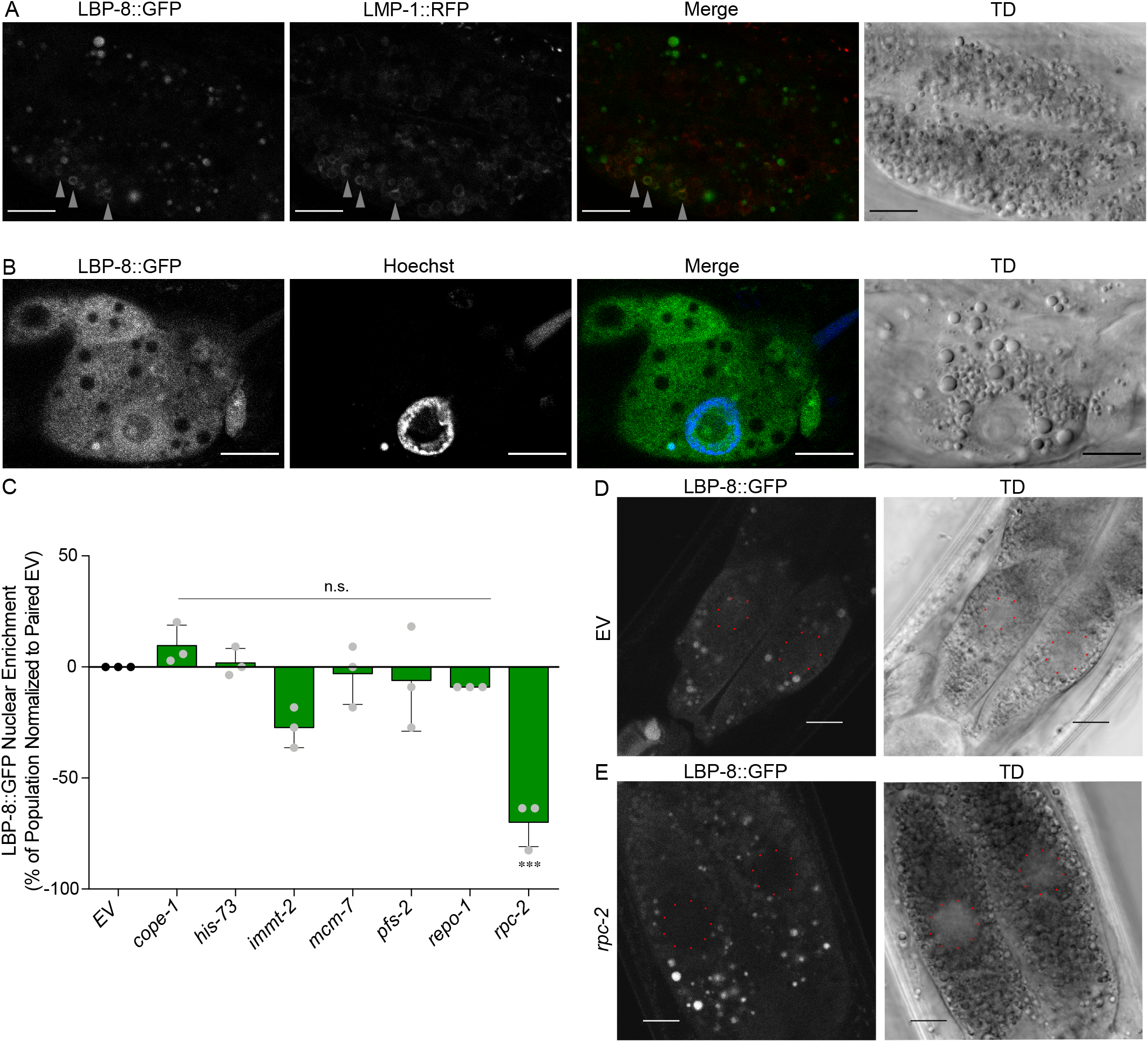
RPC-2 is Required for LBP-8 Nuclear Enrichment. (A) LBP-8::GFP co-localizes with the lysosomal marker LMP-1::RFP. Co-localization is denoted with grey triangles. (B) LBP-8::GFP co-localizes with the DNA marker Hoechst 33342. (C) LBP-8::GFP nuclear localization is reduced upon the RNAi inactivation of *rpc-2* but not *cope-1, his-73, immt-2, mcm-7, pfs-2*, or *repo-1*. Error bars represent mean +/-SD. *\*\*\*p<0*.*001*, One-Way ANOVA with Dunnett’s Post-Hoc. (D) Representative images of LBP-8::GFP showing nuclear enrichment on the EV control. (E) Representative images of LBP-8::GFP lacking nuclear enrichment upon *rpc-2* RNAi. All scale bars are 10µm.

We then inactivated the lifespan-suppressing candidates by RNAi in this transgenic strain, and examined changes in the nuclear enrichment of the LBP-8::GFP fusion. In the EV controls, we found that 72-100% of worms exhibit LBP-8 nuclear enrichment in the first pair of intestinal cells (Figure 3C). Among the lifespan-suppressing candidates, only *rpc-2* significantly affects the percentage of worms with nuclear-enriched LBP-8::GFP (*p<0*.*05*, Figure 3C). Its RNAi inactivation reduces the nuclear enrichment, and often causes LBP-8::GFP to be excluded from the nucleus (Figure 3D, 3E).

Together, these results suggest that RPC-2 binds to the LBP-8 protein and mediates its nuclear retention. RPC-2 is the homolog of RPC2, a core component of RNA Polymerase III (20) that iis responsible for transcribing tRNAs, 5S rRNAs, small nucleolar RNAs (snoRNAs) and other small RNAs (21,22). Interestingly, both the RNA polymerase III complex and the importin IMB-1 interact with the nuclear pore proteins NPP-13 and NPP-16 in *C. elegans* (23). The nuclear pore complex and importin both regulate nuclear transport of proteins and RNAs. Thus, RPC-2 might influence the nuclear-cytoplasmic distribution of LBP-8 via mediating its interaction with the nuclear pore complex and importin. It will be interesting for future studies to investigate whether RPC-2 uniquely controls the nuclear translocation of LBP-8, or its role can be generalized to other nuclear-cytoplasmic shuttling proteins.

### Nuclear import machinery is required for LBP-8 to promote longevity

Given the nuclear localization of LBP-8 and the importance of nuclear proteins for its longevity effects, we further asked whether the nuclear transportation system is involved in mediating the LBP-8 regulation of lifespan. To this end, we have conducted an RNAi screen of 14 genes encoding key components in the nuclear transportation machinery (Supplementary Table 3) (24). Using the same criteria for screening the LBP-8 binding partners, we have assessed the survival rate of the worms at day 17 of adulthood using both the *LBP-8::3XFLAG Tg* and WT worms. From the screen, we have identified two categories of candidates: one decreasing only the survival rate of the *LBP-8::3XFLAG Tg* strain but not WT and the other one decreasing both, which consist of *ima-3, imb-2, ran-1, ran-3, ran-4, xpo-1*, and *xpo-2* (Figure 4A, Table 2).

**Table 2:** Summary of Import/Export Proteins in the RNAi Screen.

| Category | *C. elegans* Gene | Human Homolog(s) | Mini Description of Functionality | WT Survival | *Tg* Survival |
| --- | --- | --- | --- | --- | --- |
| Nuclear Import | *ima-1* | KPNA1 | Importin Alpha - Adaptor for Nuclear Import Receptors |  |  |
|  | *ima-2* | KPNA1/5/6 | Importin Alpha - Adaptor for Nuclear Import Receptors |  |  |
|  | *ima-3* | KPNA4 | Importin Alpha - Adaptor for Nuclear Import Receptors |  |  |
|  | *imb-1* | KPNB1 | Importin Beta - Nuclear Import Receptor |  |  |
|  | *imb-2* | TNPO1 | Importin Beta - Nuclear Import Receptor |  |  |
|  | *imb-3* | IPO5 | Importin Beta - Nuclear Import Receptor |  |  |
|  | *xpo-2* | CSE1L | Exportin - Exporter of Importin Alpha |  |  |
| Nuclear Export | *npp-6* | NUP160 | Component of Nuclear Pore Involved in RNA Export |  |  |
|  | *npp-15* | NUP133 | Component of Nuclear Pore Involved in RNA Export |  |  |
|  | *xpo-1* | XPO1 | Exportin - Exporter of miRNA and Proteins |  |  |
|  | *xpo-3* | XPOT | Exportin - Exporter of tRNA |  |  |
| Ran Cycle | *ran-1* | RAN | RanGTPase Involved in Nuclear Import and Export |  |  |
|  | *ran-2* | RANGAP | RanGAP - Converts RanGTP to RanGDP |  |  |
|  | *ran-3* | RCC1 | RanGEF - Converts RanGDP to RanGTP |  |  |
|  | *ran-4* | NUTF2 | Imports RanGDP into the Nucleus |  |  |
|  | *ran-5* | RANBP3 | Dissociates RanGTP from Exportins in the Cytoplasm |  |  |

Note:
1. Factors in the nuclear import/export pathway are classified into “Category”. 2. Yellow marks genes whose RNAi inactivation increases survival by at least 20%, blue marks the ones decreasing survival by at least 20%, and grey marks the ones with changes less than 20%. The ones with no viable RNAi clone in the Ahringer/Vidal libraries are left blank, with *imb-1* being unavailable and *npp-6* failing to grow.

**Figure 4:**
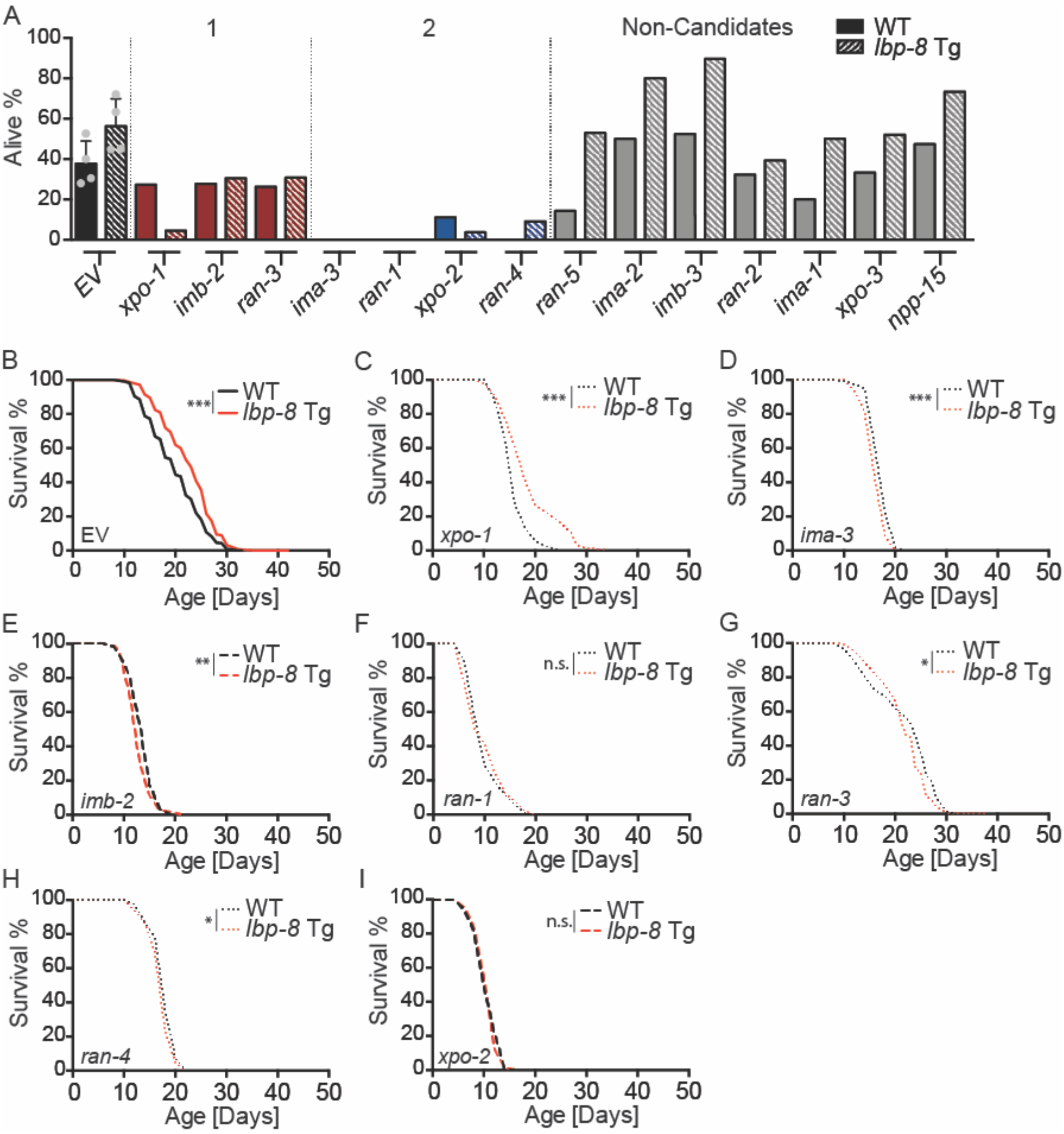
Nuclear Import but not Export is Required for LBP-8 Induced Longevity. (A) Summary of nuclear transport factors in regulating longevity: Group 1 (Red) decreasing the survival of *LBP-8::3xFLAG Tg* but not WT; Group 2 (Blue) decreasing the survival of both *LBP-8::3xFLAG Tg* and WT. Screening cutoff, >20% change in survival at day-17 adulthood. (B-I) Longitudinal analysis of the *LBP-8::3XFLAG Tg* and WT worms with EV or RNAi inactivation. The *LBP-8::3XFLAG Tg* displays lifespan extension on the EV control (B). The *LBP-8::3XFLAG Tg* cannot extend lifespan with the RNAi inactivation of *imb-2* (D) or *xpo-2* (*I*) starting at the L1 stage; or with the RNAi inactivation of *ima-3* (C), *ran-1* (E), *ran-3* (F), or *ran-4* (G) starting the L4 stage. The RNAi inactivation of *xpo-1* starting at the L4 stage does not affect the lifespan extension conferred by *LBP-8::3XFLAG Tg* (H). *\*p<0*.*05, **p<0*.*01, **p<0*.*001 Tg vs. WT* by a log-rank test.

We next used the longitudinal assay without FUDR to validate these candidates. We found that RNAi inactivation of *xpo-1* shortens the lifespans of both the *LBP-8::3XFLAG Tg* and WT worms to the same extent (Figure 4C). Thus, with the *xpo-1* knockdown, *lbp-8* is still sufficient to promote longevity. However, RNAi inactivation of all the other candidates fully abrogates the lifespan extension in the *LBP-8::3XFLAG Tg* worms (Figure 4D-I). *ima-3* and *imb-2* encode Importin-*α* and β, respectively, while *xpo-2* encodes Exportin-2, an export receptor for Importin-*α*. *ran-1* encodes the GTP-binding nuclear protein Ran, while *ran-3* and *ran-4* encode Ran GEF RCC1 and RAN-GDP importing protein NUTF2, respectively. These proteins are crucial for nuclear import of cargo proteins from the cytoplasm. On the other hand, *xpo-1* encodes Exportin-1 that mediates the nuclear export of proteins into the cytoplasm. Together, these results suggest that nuclear import but not export is specifically responsible for the longevity effect conferred by LBP-8. This finding is interesting, because the LBP-8 protein is 16.1 kDa in size, small enough to potentially diffuse through the nuclear pores on its own without the help of active nuclear import. One possibility is that LBP-8 can form a dimer to exert its longevity effect.

Mammalian FABP4 has displayed dimerization in the presence of nuclear enrichment-inducing ligands, leading to the exposure of NLS (25). Another possibility is that LBP-8 forms a complex with other proteins, which is imported into the nucleus together. One such candidate could be RPC-2, the core component of RNA polymerase III that can form a complex with the nuclear pore proteins and importin in *C. elegans* (23) and can also act as a cytosolic DNA sensor in mammalian cells (26,27).

In summary, these results reveal that only nuclear import, but not export factors are required for the pro-longevity effect caused by LBP-8. Based on this requirement, LBP-8 may function more similarly to CRABP-II, which utilizes importins but not canonical export (28), as opposed to FABP-4, which utilizes both importins and exportins (5) or FABP1/FABP2, which does not use importins and instead achieves nuclear enrichment via lessened nuclear export (29). We also discovered RPC-2 as a key regulator of LBP-8 nuclear translocation and its pro-longevity effect. This interaction between the RNA polymerase III component and FABP has not been reported before, which may provide a new way in understanding the nuclear function of FABPs. In our studies, we have focused on the nuclear but not lysosomal translocation of LBP-8, because first, there is no lysosomal proteins identified as LBP-8 binding partners through proteomic profiling; second, intrinsic autofluorescence signals derived from lysosome-like organelles in the intestine interfere with the fluorescence signal from the LBP-8::GFP fusion at the lysosomes during quantitative analyses. As a result, we currently could not determine the requirement of those binding partners in regulating LBP-8 lysosomal localization.

## Materials and Methods

### Worm Maintenance

All worms were kept at 20°C on OP50-seeded NGM plates unless stated otherwise. Worms were bleach synchronized once they reached adulthood and before the OP50 was completely consumed (starved). Worms were maintained in an unstarved state for at least 2 generations prior to any experiment being performed.

### Bleach Synchronization

Worms were synchronized using a bleaching solution. Gravid day 1 adult worms fed OP50 were washed off of NGM plates into 15mL falcon tubes using M9 media (22mM KH_2_PO_4_, 22mM Na_2_HPO_4_, 85mM NaCl, 2mM MgSO_4_). The worms were spun in a tabletop centrifuge at 3,000 rpm for 30s. The supernatant was aspirated and 10mL M9 was added. The worms were spun at 3,000 rpm for 30s, the M9 was aspirated down to 4mL, and 2mL of a freshly prepared bleaching solution (60% NaOCl, 1.6M NaOH, 1.76 mM KH_2_PO_4_, 1.76mM Na_2_HPO_4_, 6.8mM NaCl, 0.16mM MgSO_4_) was added. The falcon tube was shaken until the worms appeared paralyzed (∼30-60s). The worms were then spun at 3,000 rpm for 30s, the supernatant was aspirated, 4mL of M9 was added, followed by an additional 2mL of bleaching solution. The tubes were shaken until the adult worms were dissolved and the eggs in solution (∼30-60s). The tubes were spun at 3,000 rpm for 30s, supernatant was aspirated, and 10mL of fresh M9 added. The tubes were spun at 3,000 rpm for 30s, supernatant aspirated, and the eggs were resuspended in 4mL of M9. The eggs were left to hatch overnight while gently rocking at 20°C. The worms were plated within 48 hours of bleach synchronization.

#### Immunoprecipitation and Mass Spectrometry

Approximately 180,000 total worms were grown on 6x 15cm NGM plates seeded with 9mL of 4x concentrated OP50 to Day 1 of adulthood. Worms were washed off plates using M9 and washed twice more with M9. As much liquid was removed and worms were washed once in cold, filtered KPBS (136mM KCl and 10mM KH_2_PO_4_). The worms were resuspended in cold, filtered KPBS with 1 tablet protease inhibitor per 10mL of KPBS (cOmplete Protease Inhibitor Cocktail – Sigma 11697498001) were transferred to a cold Dounce homogenizer on ice in 2mL of KPBS. Worms were ground using approximately 100 strokes in 10 stroke increments, with a break between each increment to check a small aliquot of the worms under a dissection scope to determine if the worms’ cuticle and cells were homogenized. Once homogenized, the sample was transferred to a 2mL centrifuge tube and centrifuged at 1,000g for 3 minutes at 4°C. The supernatant was removed and combined with 80uL prewashed (3x in cold KPBS) anti-flag magnetic beads (Sigma M8823). The bead-lysate mixture was rotated at 20°C for 6 minutes. At 4°C, the supernatant was removed, and beads were gently washed 4 times with 2mL cold KPBS with protease inhibitor. The remaining liquid was removed, and beads were stored at - 80°C until mass spectrometry was performed.

### Mass Spectrometry Analysis

The beads were boiled with 2X SDS-PAGE sample buffer and subjected on 4-12% gradient gel and stained by Coomassie Brilliant Blue. The visualized bands were dissected into two parts based on protein size and destined, then in-gel digested with 500ng of MS grade trypsin (GenDepot, T9600) in lysis 50mM Ammonium bicarbonate for overnight. Tryptic peptides were extracted from gel pieces using acetonitrile and dried in a vacuum concentrator. Dried samples were resuspended in 5% methanol, 0.1% formic acid in water analyzed on Q Exactive Plus mass spectrometers (Thermo Fisher Scientific) coupled with an Easy-nLC 1000 nanoflow LC system (Thermo Fisher Scientific). An in-housed trap column packed with 1.9 μm Reprosil-Pur Basic C18 beads (2 cm × 100 μm) and separated on in-housed 5 cm × 150 μm capillary column packed with 1.9 μm Reprosil-Pur Basic C18 beads was used. A 75-min discontinuous gradient cof 4–26% acetonitrile, 0.1% formic acid at a flow rate of 800 nl/min was applied to the column then electro-sprayed into the mass spectrometer. The instrument was operated under the control of Xcalibur software version 4.0 (Thermo Fisher Scientific) in data-dependent mode, acquiring fragmentation spectra of the top 35 strongest ions. Parent mass spectrum was acquired in the Orbitrap with a full MS range of 375–1300 m/z at the resolution of 140,000. Higher-energy collisional dissociation (HCD) fragmented MS/MS spectrum was acquired in orbitrap with 17,500 resolutions. The MS/MS spectra were searched against the target-decoy C.elegans RefSeq database (release Jan. 2019, containing 28,293 entries) in Proteome Discoverer 2.1 interface (Thermo Fisher) with Mascot algorithm (Mascot 2.4, Matrix Science). The precursor mass tolerance of 20 ppm and fragment mass tolerance of 0.02Da was allowed. Two maximum missed cleavage, and dynamic modifications of acetylation of N-term and oxidation of methionine were allowed. Assigned peptides were filtered with a 1% false discovery rate (FDR) using Percolator validation based on q-value. The Peptide Spectrum Matches (PSMs) output from PD2.1 was used to group peptides onto gene level using ‘gpGrouper’ algorithm (30). This in-housed program uses a universal peptide grouping logic to accurately allocate and provide MS1 based quantification across multiple gene products. Gene-protein products (GPs) quantification was performed using the label-free, intensity-based absolute quantification (iBAQ).

### Lifespan

Bleach synchronized L1 worms were seeded on the plates and kept at 20°C throughout their entire lifespan. Worms were transferred every other day starting at the L4 stage. Movement of worms in response to a gentle stroke of the worm was used to determine if the worms was alive or dead. Individual worms were censored if they exploded (intestine protruding out of the vulva), suicided (crawled onto the plastic side of the plate and dried out), bagged (progeny hatched inside the adult), or were missing.

For lifespan assays using RNAi, RNAi was started at the L1 or L4 stage. In the event L1 feeding of the RNAi caused worms to arrest development in a larval stage, appear thin and sickly, or have a censor rate of more than 30% of the worms by day 8 of adulthood, then the lifespan was restarted with the worms being seeded on L4440 (Empty Vector (EV)) at the L1 stage and transferred to the RNAi condition at the L4 stage. RNAi clones from the laboratories of Dr. Ahringer and Dr. Vidal were sequence verified to target the gene of interest before use. The bacteria-seeded plates were left at room temperature to induce dsRNA expression in bacteria for 24 hours before use.

### RNAi Lifespan Screen

RNAi clones from the laboratories of Dr. Ahringer and Dr. Vidal and 12 well RNAi plates were used. Bleach synchronized L1 worms were seeded on the RNAi plates and kept at 20°C throughout their entire lifespan. At the L4 stage, 5-fluoro-2’deoxyuridine (FUDR) (100µM final concentration) was added to each well of the plates and the worms were counted. Starting at day 10 of adulthood, worms on the EV control were assessed to be alive or dead by tapping the 12 well plates and counting the number of worms that moved. When the *lbp-8::3xFlag* Tg worms on the EV control were at approximately 50% survival (day 17 of adulthood), all RNAi conditions for both WT and lbp-8::3xFlag Tg worms were determined to be alive or dead based on movement after 1mL of ddH2O was added to each well and the plates were gently swirled.

### Confocal Imaging

Adult worms were mounted on glass slides with 2% agarose pads and 5% NaN_3_ as a paralyzing agent. The worms were imaged on a confocal laser scanning microscope (FV3000, Olympus).

### Nuclear Enrichment Imaging and Analysis

Bleach synchronized L1 worms were seeded on plates with HT115 bacteria. The same seeding scheme was used for this as for the Lifespans, where RNAi was started at either the L1 or L4 stage depending upon the RNAi’s effect on development, perceived health, or censor rate. If RNAi was started at the L1 stage, then Day 1 adults were analyzed, but if RNAi was started at the L4 stage, then Day 2 adults were analyzed.

Approximately 20 adult worms were picked into a 6µL droplet of 5% NaN_3_ on a square glass coverslip. The worms were gently separated once they were paralyzed and the coverslip was mounted onto a 2% agarose pad on a glass slide. The worms were then imaged on a confocal laser scanning microscope (FV3000, Olympus).

The first pair of intestinal cells were imaged. First, at least one of the nuclei of the first pair was set into focus using the TD channel. An image was then taken using both the TD and GFP channels. If nuclear enrichment was not observed in the initial image, then the GFP channel was used to slowly ascend and descend along the Z axis while continuously scanning. If an approximately nucleus-sized region with increased fluorescence relative to the surrounding area appeared on another focal plane, then another image was taken using both the GFP and TD channels.

To analyze the nuclear enrichment, FIJI was used to analyze the images taken on the FV3000. Briefly, the images of one replicate of one condition were simultaneously opened in FIJI, and it was ensured that only the GFP channel was visible. Using the GFP channel’s image, if there was a perceived increase in signal relative to the adjacent cytoplasm, then the TD channel was used to confirm if this increase in signal is the nucleus. Images were scored as having either “Enrichment” or “No Enrichment” relative to the cytoplasm. If two images were taken for a single worm, then only one score was used, with Enrichment trumping No Enrichment.

### Hoechst Staining

100µL of 1:10 dilution of Hoechst 33342 (H3570, Thermofisher) was added to the plate and kept in the dark at room temperature for 24 hours before being used. L4 stage worms were transferred to the Hoechst plates and kept in the dark at 20°C. 24 hours later, the adult worms were mounted in 5% NaN_3_ on a 2% agarose pad on a glass slide and imaged on a confocal laser scanning microscope (FV3000, Olympus).

## Acknowledgements

We would like to thank A. Dervisefendic and P. Svay for maintenance support. This work has been supported by the NIH grants R01AG045183 (M.C.W.), R01AT009050 (M.C.W.), R01AG062257 (M.C.W.), DP1DK113644 (M.C.W.), Welch Foundation Q-1912 (M.C.W.). M.C.W is a Howard Hughes Medical Institute Investigator.

---

**Supplemental Table 1: iBAQ Scores, Normalization, and Fold Changes of LBP-8 IP/MS**

**Supplemental Table 2: Survival Data from Lifespan Screen**

**Supplemental Table 3: Lifespan Tables**

